# Global gene expression analysis of *Streptococcus agalactiae* at exponential growth phase

**DOI:** 10.1101/2020.11.13.381939

**Authors:** Inês Silvestre, Vítor Borges, Sílvia Duarte, Alexandra Nunes, Rita Sobral, Luís Vieira, João Paulo Gomes, Maria José Borrego

## Abstract

*Streptococcus agalactiae* is a leading cause of neonatal infections and an increasing cause of infections in adults with underlying diseases. One of the first *S. agalactiae* isolates to be subjected to whole genome sequencing was NEM316, a strain responsible for a fatal case of septicemia that has been widely used as reference strain for *in vitro* assays. Whole transcriptome analyses may provide an essential contribute to the understanding of the molecular mechanisms responsible for bacteria adaptation and pathogenicity, still, so far, very few studies were dedicated to the analysis of global gene expression of *S. agalactiae*. Here, we applied RNA-sequencing to perform a comparative overview of the global gene expression levels of the *S. agalactiae* reference strain NEM316 at the exponential growth phase. Genes were ranked by expression level and grouped by functional category and 46% of the top-100 expressed genes encode proteins involved in “Translation, ribosomal structure and biogenesis”. Among the group of highly expressed genes were also represented genes with no assigned functional category. Although this result warrants further investigation, most of them might be implicated in stress response. As very little is known about the molecular mechanisms behind the release of DNase’s *in vitro* and *in vivo*, we also performed preliminary assays to understand whether direct DNA exposure affects the gene expression of strain NEM316 at the exponential growth phase. No differentially expressed genes were detected, which indicates that follow-up studies are needed to disclose the complex molecular pathways (and stimuli) triggering the release of DNase’s. In general, we provide data on the global expression levels of NEM316 at exponential growth phase that may contribute to better understand *S. agalactiae* adaptation and virulence.

## 1. Introduction

*Streptococcus agalactiae* belongs to the family of streptococcaceae and is a common inhabitant of the healthy human gut and urogenital tract, particularly in women [1]. As an opportunistic pathogen, *S. agalactiae* is a major cause of neonatal infections and mortality, and has also emerged as a pathogen in elderly and immunocompromised adults [2]. *S. agalactiae* is capable to adhere to various host cell types, namely epithelial cells of the vagina and the lung, endothelial cells and micro-vascular endothelial cells of the blood-brain barrier [3]. However, the molecular mechanisms underlying the transition from colonization to infection have yet to be disclosed, and while the exact pathogenic features of *S. agalactiae* remain unrevealed, many virulence factors have been proposed to explain infection-related pathogenesis [4]. In *S. agalactiae* reference strain NEM316 (genotype: III/ ST23) [5], isolated from an infected infant, several extracytoplasmic virulence factors have been identified, such as capsule, proteases, adhesins, haemolysin, pili and pigment. These factors mediate adhesion and epithelial cell invasion, and/or antagonize the immune system during phagocytosis [1, 6].

Understanding the transcriptomic setting allows to obtain insights on the pathways of bacterial physiology, metabolism, and adaptation to changing environments [7, 8]. Over the past decade, RNA sequencing (RNA-seq) has become an indispensable tool for transcriptome-wide analysis of differential gene expression. Together with improved computational tools for data analysis, innovations in RNA-seq technologies are contributing to a better understanding of RNA biology and intermolecular interactions that govern RNA function, as well as transcriptional dynamics changes driving bacterial response and adaptation, to distinct growth conditions and external stimuli [9]. Recently, RNA-seq technology has been successfully applied to clarify some molecular mechanisms of pathogenesis in *S. agalactiae* [8, 10-15]. The first *S. agalactiae* comparative transcriptomic study evidenced several genetic factors (in particular related to lactose metabolism) likely important in adaptation to the bovine environment [10]. In 2018, Hooven *et al.* [12] identified the gene products necessary for *S. agalactiae* survival in human whole blood, and Cook *et al.* [13] identified novel genes involved in murine vaginal colonization by *S. agalactiae*. The contribute of CRISPR-associated protein-9 to *S. agalactiae* colonization and disease was also investigated by RNA-seq [14]. While these targeted studies provided insightful data about specific *S. agalactiae* adaptive traits, the expression levels of reference strains during their normal growth in the laboratory have not yet been systematized. In this study, we investigated the global gene expression of *S. agalactiae* NEM316 during the exponential growth phase by RNA-seq. We also performed preliminary assays to understand whether direct DNA exposure affects *S. agalactiae* gene expression.

## 2. Materials and methods

### 2.1. Whole genome sequencing of the laboratory reference strain NEM316

All experiments were conducted using the *S. agalactiae* reference strain NEM316 (ATCC 12403; genotype: III/ ST23). The reference strain NEM316, maintained in laboratory at −80°C in cryopreservation tubes (Cryoinstant Red, VWR, Belgium), was cultured in Columbia agar supplemented with 5% sheep blood (Biomérieux, Marcy l’Etoile, France) at 5% CO_2_ for 24 h. These cultures were used to inoculate cultures of fresh Todd Hewitt broth (THB) supplemented with 0,5% yeast extract that were allowed to incubate without shaking at 37°C with 5% CO_2_. Cell growth was monitored by measuring the optical density at 600 nm (OD_600_). At middle exponential phase (OD_600_=0,2-0,5), 1 ml of bacterial cells were collected by centrifugation (3000 rpm, 10 min), resuspended in 200μl of PBS and immediately stored at −20°C for further DNA extraction. Genomic DNA was extracted as previously referred [16], with minor changes. Briefly, bacterial cells were subjected to a high-speed centrifugation (14,000 rpm) for 10 min at 4°C. The pellet was digested for 2 h at 37°C with 200 μl of Tris-EDTA buffer, pH 8.0, containing 10 U mutanolysin (Sigma-Aldrich, St. Louis, USA) and 15 mg/ml lysozyme (Sigma-Aldrich, St. Louis, USA) before treatment with 10 mg/ml proteinase K (Roche, Penzberg, Germany). Subsequently, DNA was extracted using the NucliSENS® EasyMag® (BioMérieux, Marcy l’Etoile, France) according to manufacturer’s instructions. The concentration of the extracted DNA was measured with Qubit™ (ThermoFisher Scientific, Massachusetts, USA), and then subjected to Next Generation Sequencing in NextSeq 550 equipment (2×150bp) (Illumina, USA). The reads were deposited in the European Nucleotide Archive (ENA) (Bioproject PRJEB41294) under the accession number ERR4836035.

In order to evaluate the genetic differences between the genome of our laboratory passaged reference strain and the publicly available genome (GenBank accession number NC004368), two strategies were applied: i) de novo genome assembly using INNUca v.4.0.1 (https://github.com/B-UMMI/INNUca) [17], followed by genome alignment and inspection using MAUVE (http://darlinglab.org/mauve/mauve.html) (to inspect for the presence of structural changes, such as large indels); ii) reference-based mapping using Snippy v3.2 (to detect SNPs and small indels) (https://github.com/tseemann/snippy).

Clusters of Orthologous Groups (COGs) categories were assigned to the amino acid sequences retrieved from the NEM316 NCBI annotation (GenBank accession number NC004368) using “cdd2cog” script [18] after RPS-BLAST+ (Reverse Position-Specific BLAST) (e-value cut-off of 1e-2), where only the best hit (lowest e-value) and first COG were considered.

### 2.2. Bacterial culture for RNA-seq

Bacterial clones of NEM316 were grown in Columbia agar supplemented with 5% sheep blood (Biomérieux, Marcy l’Etoile, France) with 5% CO_2_ for 24 h and then inoculated in Todd Hewitt selective media broth (THB) that were allowed to incubate without shaking at 37°C, 5% CO_2_. Cell growth of *S. agalactiae* strain NEM316 was monitored by optical density at 600nm (OD_600_). At OD_600_=0,6 (exponential growth phase, see Supplementary material Figure S1) 1 ml of bacterial cells were collected by centrifugation (3000 rpm, 10 min), resuspended in 1,8 ml of fresh THB and incubated for 0, 10 and 20 minutes at 37°C in the presence of 200μl of PBS (used as control). For direct DNA exposure assays, the same procedure was performed, with the exception that 2μg/ml of DNA (human DNA extracted from Hela cells, used as a stimulus) was added instead of PBS, and nuclease reaction was stopped by adding EDTA (0.5 M, pH 8.0) at 4°C. For both conditions, 1 ml of each bacterial culture was collected and immediately subjected to high-speed centrifugation (14,000 rpm) for 10 min at 4°C for RNA extraction. Note that we intentionally did not treat the bacterial culture from which RNA would be extracted with RNAprotect™ Bacteria Reagent (Qiagen, CA, USA) because a preliminary assay showed that this product degrades *S. agalactiae* RNA (data not shown).

### 2.3. RNA extraction

RNA was extracted as previously referred [16], with minor changes. Briefly, the cells were lysed in 200 μl of Tris-EDTA buffer, pH 8.0, containing 10 U mutanolysin (Sigma-Aldrich, St. Louis, USA) and 15 mg/ml lysozyme (Sigma-Aldrich, St. Louis, USA), at 37°C during 90 min. The RNeasy mini kit (Qiagen, CA, USA) was used according to manufacturer’s instructions. Residual contaminant DNA was removed using 30 U RNase-free DNase (Qiagen CA, USA), and elution was performed with 40 μl of RNase-free water. RNA yield and purity were determined by absorbance measurement at 260 and 280 nm using the Nanodrop 1000 spectrophotometer (Thermo Fisher Scientific, Massachusetts, USA). Extracted RNA was finally stored at −80°C until use.

### 2.4. Bacterial mRNA preparation/purification

Bacterial mRNA was enriched using the Ribo-Zero™ rRNA Removal Kit (Illumina, CA, USA) which removed abundant 16S and 23S rRNA from total RNA. The obtained bacterial mRNA was concentrated to a final volume of 14 μl using the RNeasy® MinElute® Cleanup Kit (Qiagen, CA, USA). The yield and integrity of the enriched mRNA samples was assessed with an Agilent Bioanalyzer, where the absence of rRNA readings is indicative of the success of rRNA depletion and purity of mRNA.

### 2.5. RNA-seq

Bacterial mRNA-enriched samples were subjected to library construction by TruSeq Stranded mRNA sample preparation kit (Illumina, CA, USA). The obtained cDNA libraries were subjected to RNA-seq on a high-throughput MiSeq Illumina apparatus, targeting around 4M reads per 1Mbp. Sequence reads (2×75bp) were subjected to quality control and subsequently mapped to the *S. agalactiae* NEM316 genome (as obtained above) using Bowtie2 [19]. Relative gene expression was quantified and normalized as fragments per kb of CDS per million mapped reads (FPKM) using the Cufflinks software (version 2.1.1; http://cufflinks.cbcb.umd.edu/). For comparative global gene expression analyses between normal growth condition and the “direct DNA exposure”, we applied HTSeq-count (https://htseq.readthedocs.io/en/release0.11.1/count.html#) for read counting and state-of-the-art software for differential expression analysis (namely, EdgeR and Voom/Limma) using the interactive web-tool DEGUST (https://degust.erc.monash.edu/) [20].

The reads were deposited in ENA (Bioproject PRJEB41294) under the accession numbers ERR4836029, ERR4836030, ERR4836031, ERR4836032, ERR4836033, ERR4836034.

## 3. Results and discussion

### 3.1. NEM316 whole genome sequencing

*S. agalactiae* reference strain NEM316 was isolated 18 years ago [5] and, since then, has been maintained in laboratory. As such, in order to prepare RNA-seq assays, NEM316 was subjected to WGS to evaluate whether this laboratory passaged strain had significant genetic changes in comparison with the publicly available genome [5] (GenBank accession number NC004368). Six genome-dispersed mutations were detected, including three small indels and three single nucleotide polymorphisms (SNPs), corresponding to three non-synonymous mutations (Table 1). Among these, we highlight a SNP in *relA*, which encodes an enzyme known to be involved in stringent response and bacterial adaptation to environmental stress [12]. In *S. agalactiae*, *relA* knockout strains demonstrated decreased expression of β-hemolysin/cytolysin, an important cytotoxin implicated in facilitating invasion [12]. Although the impact of these particular mutations at the transcriptomic level is unknown, we cannot rule out the possibility that they reflect events of laboratory adaptation. Notwithstanding, we consider a good practice to analyze the genome backbone of strains subjected to gene expression (or other *in vitro*) assays, as a means to provide more complete data required to better interpret and discuss the results.

**Table 1.** Genomic alterations for *S. agalactiae* NEM316 maintained in laboratory.

| Mutation location <sup>a</sup> | Type | Locus tag <sup>b</sup> | Old locus tag <sup>b</sup> | nt (aa) change <sup>c</sup> | Effect | Product |
| --- | --- | --- | --- | --- | --- | --- |
| 1313729 | insertion | GBS_RS06735.<br>pseudogene | gbs1273 | 811insA<sup>d</sup> | --- | glucose-1-phosphate<br>thymidyltransferase |
| 1363940 | insertion |  |  | CTT > CTTT |  |  |
| 1741404 | deletion |  |  | GATATATA<br>> GATATA |  |  |
| 1839016 | SNP | GBS_RS09285 | gbs1779 | A1077T<br>(Glu359Asp) | Missense<br>variant | Major facilitator superfamily<br>transporter |
| 2001001 | SNP | GBS_RS10040 | gbs1928 | G1952C<br>(Trp651Ser) | Missense<br>variant | bifunctional (p)ppGpp<br>synthetase/guanosine-3',5'-<br>bis(diphosphate) 3'-<br>pyrophosphohydrolase (relA) |
| 2129967 | SNP | GBS_RS10675 | gbs2055 | A11T<br>(Lys4Met) | Missense<br>variant | arginine repressor |

<sup>a</sup> Polymorphism location refers to the location in the reference genome: NEM316 (GenBank accession number NC004368).
<sup>b</sup> Open Reading Frames (ORFs) designations according to reference genome: NEM316 (GenBank accession number NC004368).
<sup>c</sup> The nucleotide changes in open reading frames are presented in the 5' to 3' direction.
<sup>d</sup> This locus is a pseudogene in the reference genome (GenBank accession number NC004368) due to a one bp deletion as such NEM316 of our laboratory retains the original not truncated allele.

### 3.2. NEM316 transcriptomic analyses

The main goal of the present study was to evaluate the global gene expression dynamics of NEM316 *S. agalactiae* strain during the exponential phase using the RNA-seq technology. This growth phase is particularly interesting when studying the transcriptional activity because most cells in the population are actively dividing and this ensures that the expression of most of the *S. agalactiae* genes is assessed. As such, three time points (0, 10 and 20 minutes) were evaluated during the exponential growth phase in THB.

Firstly, we ranked the *S. agalactiae* genes by expression level and correlated them with the gene functional category (Figure 1) (Supplementary material Table S1). Huge differences in the median expression levels were observed between different gene functional categories (Figure 1), with the top expressed functional category (“Translation, ribosomal structure and biogenesis”) revealing a median expression value that was 55-fold higher than the less expressed functional category (“Cell motility”). The top three most expressed functional categories were “Translation, ribosomal structure and biogenesis”, “Energy production and conversion” and “Posttranslational modification, protein turnover, chaperones”. This result may not be surprising as during the exponential growth phase, the cell division rate is maximum and this implicates a high demand for proteins playing a role in translation and metabolism. The three functional categories with the lowest median expression levels were “Cell motility”, “Intracellular trafficking, secretion, and vesicular transport” and “Not assigned/Function unknown”.

**Figure 1.**
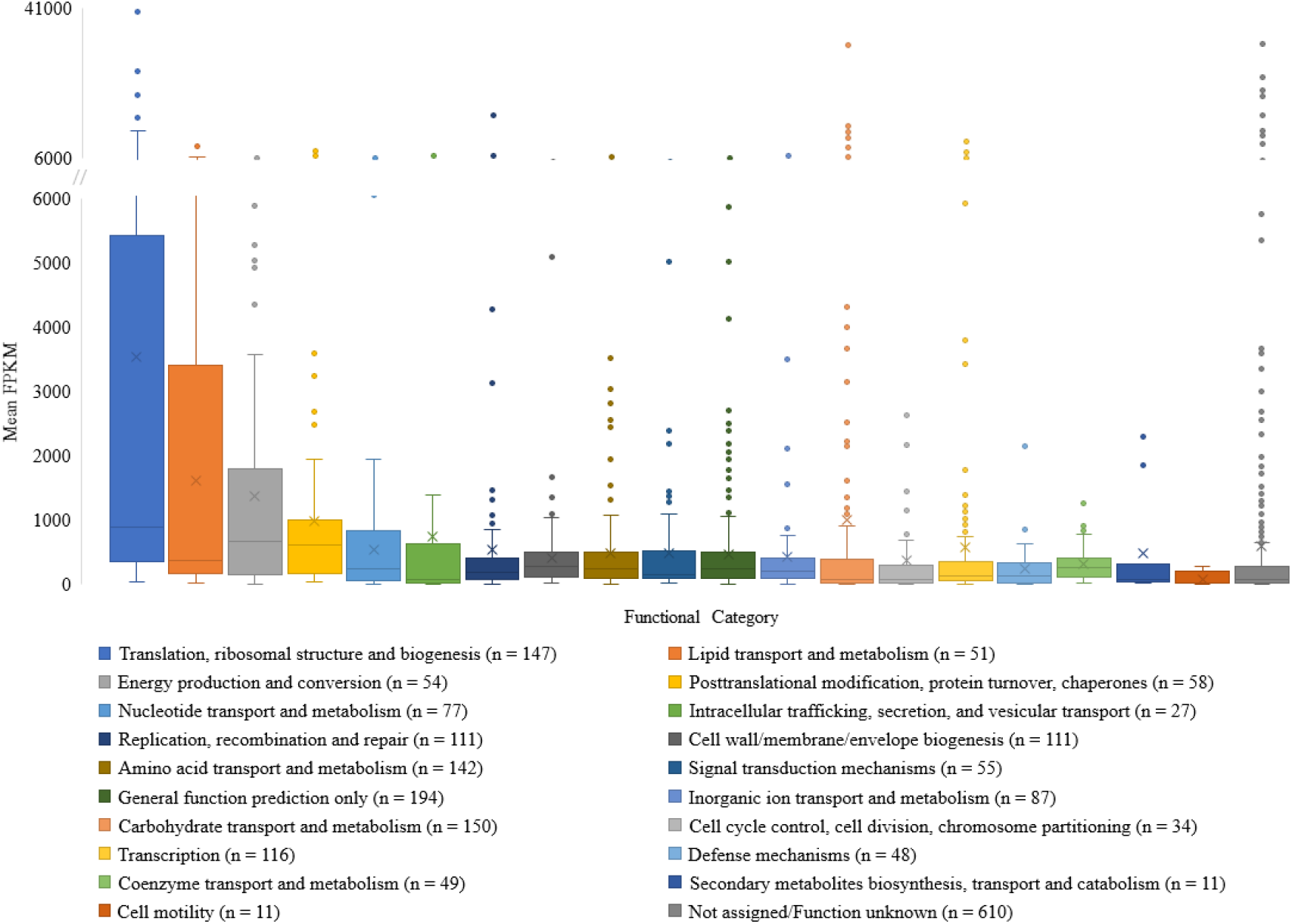
Global gene expression by functional categories. Box plot showing the distribution of gene expression levels by functional category, for reference strain NEM16 at the exponential growth phase. Values reflect the mean expression level evaluated at three time points (0, 10 and 20 min) during exponential growth. Genes with mean expression levels < 1 FPKM were excluded from the analysis.

The analysis of the top-100 genes with highest level of expression at the exponential growth phase (Table 2) showed that the genes belonging to the functional category “Translation, ribosomal structure and biogenesis” were highly represented (46%), with the proportion of genes from the remainder functional categories never exceeding 15%. Also, 15 genes with unknown function were detected among the highly expressed genes (GBS_RS11490, GBS_RS06375, GBS_RS06390, GBS_RS06380, GBS_RS06405, GBS_RS06400, GBS_RS06385, GBS_RS06800, GBS_RS11205, GBS_RS03445, GBS_RS10615, GBS_RS06395, GBS_RS00250, GBS_RS05190, GBS_RS11525). Although these genes have not been grouped into any functional category by RPS-BLAST against the COG database, fine-tune evaluation of their putative function based on the new NEM316 genome annotation (released on June 2020), plus literature surveys, provided some clues that might justify the observed high expression level. This is the case of GBS_RS00250, which is believed to be required for *S. agalactiae* cell division due to its potential role in peptidoglycan cleavage, since it includes a CHAP domain that has been associated with peptidoglycan hydrolysis [21]. Disruption of this gene was shown to cause an altered cell morphology and an increased susceptibility towards different antibiotics, namely β-lactam antibiotics [22, 23]. GBS_RS05190, a PASTA domain-containing protein, may also be involved in bacterial cell division as PASTA repeats are known to be key regulators of the membrane during bacterial cell division [24]. GBS_RS11205 is a putative holin-like toxin, and holins, which are encoded by phages, have been considered responsible for disruption of the cytoplasmic membrane to assist endolysins during cell lysis [25]. Although most of these proteins are implicated in the bacterial stress response, like GBS_RS06375, GBS_RS06380, GBS_RS06405, GBS_RS06400, GBS_RS06385, GBS_RS03445 and GBS_RS06395 [10, 26-28], others (GBS_RS11490, GBS_RS06390, GBS_RS06800, GBS_RS10615, GBS_RS11525) do not have any assigned function, neither any predicted functional domain.

The functional category “Intracellular trafficking, secretion, and vesicular transport”, which belongs to one of the functional categories with the lowest median expression levels, was also represented among the top-100 most expressed genes by GBS_RS09920 and GBS_RS00560 (preprotein translocase subunit YajC and preprotein translocase subunit SecY, respectively).

**Table 2.** Top-100 most expressed genes by functional category.

| Locus tag <sup>a</sup> | Old locus tag <sup>a</sup> | Product <sup>b</sup> | Functional Category <sup>c</sup> | Mean FPKM <sup>d</sup> |
| --- | --- | --- | --- | --- |
| GBS_RS04310 | gbs0782 | elongation_factor_Tu | Translation, ribosomal structure and biogenesis | 40081 |
| GBS_RS11490 | --- | Not assigned | Not assigned/Function unknown | 36837 |
| GBS_RS06375 | gbs1202 | Asp23/Gls24_family_envelope_stress_response_protein | Not assigned/Function unknown | 32678 |
| GBS_RS09445 | gbs1811 | type_I_glyceraldehyde-3-phosphate_dehydrogenase | Carbohydrate transport and metabolism | 32335 |
| GBS_RS09575 | gbs1836 | 50S_ribosomal_protein_L34 | Translation, ribosomal structure and biogenesis | 26121 |
| GBS_RS06390 | gbs1205 | DUF2273_domain-containing_protein | Not assigned/Function unknown | 24824 |
| GBS_RS06380 | gbs1203 | CsbD_family_protein | Not assigned/Function unknown | 22768 |
| GBS_RS06405 | gbs1208 | GlsB/YeaQ/YmgE_family_stress_response_membrane_protein | Not assigned/Function unknown | 21825 |
| GBS_RS11120 | --- | 50S_ribosomal_protein_L36 | Translation, ribosomal structure and biogenesis | 20731 |
| GBS_RS06400 | gbs1207 | GlsB/YeaQ/YmgE_family_stress_response_membrane_protein | Not assigned/Function unknown | 20358 |
| GBS_RS03205 | --- | HU_family_DNA-binding_protein | Replication, recombination and repair | 15942 |
| GBS_RS06385 | gbs1204 | Asp23/Gls24_family_envelope_stress_response_protein | Not assigned/Function unknown | 15927 |
| GBS_RS00885 | --- | 50S_ribosomal_protein_L28 | Translation, ribosomal structure and biogenesis | 15451 |
| GBS_RS00525 | gbs0071 | type_Z_30S_ribosomal_protein_S14 | Translation, ribosomal structure and biogenesis | 15276 |
| GBS_RS03485 | gbs0608 | phosphopyruvate_hydratase | Carbohydrate transport and metabolism | 13436 |
| GBS_RS06800 | --- | DUF3042_family_protein | Not assigned/Function unknown | 12964 |
| GBS_RS00875 | gbs0125 | fructose-bisphosphate_aldolase | Carbohydrate transport and metabolism | 12483 |
| GBS_RS11205 | --- | putative_holin-like_toxin | Not assigned/Function unknown | 12444 |
| GBS_RS01260 | --- | 30S_ribosomal_protein_S15 | Translation, ribosomal structure and biogenesis | 12269 |
| GBS_RS04595 | gbs0839 | phosphocarrier_protein_HPr | Carbohydrate transport and metabolism | 12076 |
| GBS_RS03445 | --- | CsbD_family_protein | Not assigned/Function unknown | 11868 |
| GBS_RS00545 | gbs0075 | 30S_ribosomal_protein_S5 | Translation, ribosomal structure and biogenesis | 11614 |
| GBS_RS03385 | gbs0587 | 50S_ribosomal_protein_L19 | Translation, ribosomal structure and biogenesis | 11587 |
| GBS_RS00495 | gbs0065 | 50S_ribosomal_protein_L16 | Translation, ribosomal structure and biogenesis | 11233 |
| GBS_RS10615 | gbs2043 | DUF1292_domain-containing_protein | Not assigned/Function unknown | 11174 |
| GBS_RS00510 | gbs0068 | 50S_ribosomal_protein_L14 | Translation, ribosomal structure and biogenesis | 11111 |
| GBS_RS00570 | gbs0080 | translation_initiation_factor_IF-1 | Translation, ribosomal structure and biogenesis | 10915 |
| GBS_RS09435 | gbs1809 | phosphoglycerate_kinase | Carbohydrate transport and metabolism | 10578 |
| GBS_RS00535 | gbs0073 | 50S_ribosomal_protein_L6 | Translation, ribosomal structure and biogenesis | 10415 |
| GBS_RS07230 | gbs1374 | 50S_ribosomal_protein_L7/L12 | Translation, ribosomal structure and biogenesis | 10230 |
| GBS_RS00515 | gbs0069 | 50S_ribosomal_protein_L24 | Translation, ribosomal structure and biogenesis | 10060 |
| GBS_RS10665 | gbs2053 | cold-shock_protein | Transcription | 9839 |
| GBS_RS00505 | gbs0067 | 30S_ribosomal_protein_S17 | Translation, ribosomal structure and biogenesis | 9523 |
| GBS_RS00540 | gbs0074 | 50S_ribosomal_protein_L18 | Translation, ribosomal structure and biogenesis | 9357 |
| GBS_RS00580 | gbs0083 | 30S_ribosomal_protein_S11 | Translation, ribosomal structure and biogenesis | 9271 |
| GBS_RS05090 | --- | 30S_ribosomal_protein_S20 | Translation, ribosomal structure and biogenesis | 9175 |
| GBS_RS06395 | gbs1206 | alkaline_shock_response_membrane_anchor_protein_AmaP | Not assigned/Function unknown | 9153 |
| GBS_RS00575 | gbs0082 | 30S_ribosomal_protein_S13 | Translation, ribosomal structure and biogenesis | 8859 |
| GBS_RS02085 | gbs0337 | acetyl-CoA_carboxylase_biotin_carboxyl_carrier_protein | Lipid transport and metabolism | 8621 |
| GBS_RS00585 | gbs0084 | DNA-directed_RNA_polymerase_subunit_alpha | Transcription | 8452 |
| GBS_RS04600 | gbs0840 | phosphoenolpyruvate--protein_phosphotransferase | Carbohydrate transport and metabolism | 8379 |
| GBS_RS00490 | gbs0064 | 30S_ribosomal_protein_S3 | Translation, ribosomal structure and biogenesis | 8315 |
| GBS_RS03355 | --- | type_B_50S_ribosomal_protein_L31 | Translation, ribosomal structure and biogenesis | 8098 |
| GBS_RS07870 | --- | 30S_ribosomal_protein_S21 | Translation, ribosomal structure and biogenesis | 8026 |
| GBS_RS09175 | gbs1757 | 30S_ribosomal_protein_S18 | Translation, ribosomal structure and biogenesis | 7951 |
| GBS_RS00555 | gbs0077 | 50S_ribosomal_protein_L15 | Translation, ribosomal structure and biogenesis | 7785 |
| GBS_RS09765 | gbs1874 | peroxiredoxin | Posttranslational modification, protein turnover, chaperones | 7672 |
| GBS_RS00590 | --- | 50S_ribosomal_protein_L17 | Translation, ribosomal structure and biogenesis | 7638 |
| GBS_RS00475 | gbs0061 | 50S_ribosomal_protein_L2 | Translation, ribosomal structure and biogenesis | 7591 |
| GBS_RS07235 | gbs1375 | 50S_ribosomal_protein_L10 | Translation, ribosomal structure and biogenesis | 7576 |
| GBS_RS00520 | gbs0070 | 50S_ribosomal_protein_L5 | Translation, ribosomal structure and biogenesis | 7457 |
| GBS_RS04180 | gbs0756 | PspC_domain-containing_protein | Transcription | 7382 |
| GBS_RS04315 | gbs0783 | triose-phosphate_isomerase | Carbohydrate transport and metabolism | 6956 |
| GBS_RS00460 | gbs0058 | 50S_ribosomal_protein_L3 | Translation, ribosomal structure and biogenesis | 6902 |
| GBS_RS09450 | gbs1812 | elongation_factor_G | Translation, ribosomal structure and biogenesis | 6790 |
| GBS_RS04440 | gbs0808 | superoxide_dismutase | Inorganic ion transport and metabolism | 6587 |
| GBS_RS09920 | gbs1904 | preprotein_translocase_subunit_YajC | Intracellular trafficking, secretion, and vesicular transport | 6572 |
| GBS_RS09180 | gbs1758 | single-stranded_DNA-binding_protein | Replication, recombination and repair | 6519 |
| GBS_RS09770 | gbs1875 | alkyl_hydroperoxide_reductase_subunit_F | Posttranslational modification, protein turnover, chaperones | 6456 |
| GBS_RS00560 | gbs0078 | preprotein_translocase_subunit_SecY | Intracellular trafficking, secretion, and vesicular transport | 6382 |
| GBS_RS02055 | gbs0331 | ketoacyl-ACP_synthase_III | Lipid transport and metabolism | 6364 |
| GBS_RS02105 | gbs0341 | acetyl-CoA_carboxylase_carboxyl_transferase_subunit_alpha | Lipid transport and metabolism | 6312 |
| GBS_RS00480 | gbs0062 | 30S_ribosomal_protein_S19 | Translation, ribosomal structure and biogenesis | 6299 |
| GBS_RS04320 | gbs0784 | phosphoglycerate_mutase | Carbohydrate transport and metabolism | 6283 |
| GBS_RS08050 | gbs1534 | amino_acid_ABC_transporter_ATP-binding_protein | Amino acid transport and metabolism | 6171 |
| GBS_RS02095 | gbs0339 | acetyl-CoA_carboxylase_biotin_carboxylase_subunit | Lipid transport and metabolism | 6123 |
| GBS_RS00465 | gbs0059 | 50S_ribosomal_protein_L4 | Translation, ribosomal structure and biogenesis | 6070 |
| GBS_RS04980 | gbs0916 | nucleoside-diphosphate_kinase | Nucleotide transport and metabolism | 6062 |
| GBS_RS01325 | gbs0210 | 30S_ribosomal_protein_S9 | Translation, ribosomal structure and biogenesis | 5995 |
| GBS_RS07565 | gbs1438 | 50S_ribosomal_protein_L27 | Translation, ribosomal structure and biogenesis | 5926 |
| GBS_RS02050 | gbs0330 | MarR_family_transcriptional_regulator | Transcription | 5924 |
| GBS_RS04805 | gbs0882 | F0F1_ATP_synthase_subunit_epsilon | Energy production and conversion | 5896 |
| GBS_RS05130 | gbs0946 | FAD-dependent_oxidoreductase | General function prediction only | 5876 |
| GBS_RS00250 | gbs0016 | CHAP_domain-containing_protein | Not assigned/Function unknown | 5757 |
| GBS_RS00455 | gbs0057 | 30S_ribosomal_protein_S10 | Translation, ribosomal structure and biogenesis | 5733 |
| GBS_RS02100 | gbs0340 | acetyl-CoA_carboxylase_carboxyltransferase_subunit_beta | Lipid transport and metabolism | 5565 |
| GBS_RS09455 | --- | 30S_ribosomal_protein_S7 | Translation, ribosomal structure and biogenesis | 5501 |
| GBS_RS00485 | gbs0063 | 50S_ribosomal_protein_L22 | Translation, ribosomal structure and biogenesis | 5480 |
| GBS_RS09760 | gbs1873 | 30S_ribosomal_protein_S2 | Translation, ribosomal structure and biogenesis | 5366 |
| GBS_RS05190 | gbs0958 | PASTA_domain-containing_protein | Not assigned/Function unknown | 5360 |
| GBS_RS02005 | gbs0326 | ribosome-associated_translation_inhibitor_RaiA | Translation, ribosomal structure and biogenesis | 5352 |
| GBS_RS04875 | gbs0895 | thiamine_pyrophosphate-dependent_dehydrogenase_E1_component_subunit_alpha | Energy production and conversion | 5285 |
| GBS_RS02080 | gbs0336 | beta-ketoacyl-ACP_synthase_II | Lipid transport and metabolism | 5201 |
| GBS_RS07760 | gbs1477 | PI-2a_pilus_major_subunit_PilB | Cell wall/membrane/envelope biogenesis | 5099 |
| GBS_RS10710 | gbs2062 | 50S_ribosomal_protein_L32 | Translation, ribosomal structure and biogenesis | 5094 |
| GBS_RS03295 | gbs0569 | (S)-acetoin_forming_diacetyl_reductase | Lipid transport and metabolism | 5079 |
| GBS_RS02045 | gbs0329 | enoyl-CoA_hydratase | Lipid transport and metabolism | 5075 |
| GBS_RS00470 | gbs0060 | 50S_ribosomal_protein_L23 | Translation, ribosomal structure and biogenesis | 5075 |
| GBS_RS04800 | gbs0881 | F0F1_ATP_synthase_subunit_beta | Energy production and conversion | 5029 |
| GBS_RS08755 | gbs1672 | response_regulator_transcription_factor | Signal transduction mechanisms | 5021 |
| GBS_RS07510 | gbs1427 | KH_domain-containing_protein | General function prediction only | 5012 |
| GBS_RS11525 | --- | Not assigned | Not assigned/Function unknown | 4975 |
| GBS_RS07575 | gbs1440 | 50S_ribosomal_protein_L21 | Translation, ribosomal structure and biogenesis | 4954 |
| GBS_RS05135 | gbs0947 | L-lactate_dehydrogenase | Energy production and conversion | 4926 |
| GBS_RS09460 | gbs1814 | 30S_ribosomal_protein_S12 | Translation, ribosomal structure and biogenesis | 4883 |
| GBS_RS02060 | gbs0332 | acyl_carrier_protein | Lipid transport and metabolism | 4739 |
| GBS_RS01320 | gbs0209 | 50S_ribosomal_protein_L13 | Translation, ribosomal structure and biogenesis | 4733 |
| GBS_RS09380 | gbs1798 | 30S_ribosomal_protein_S14 | Translation, ribosomal structure and biogenesis | 4605 |
| GBS_RS00530 | gbs0072 | 30S_ribosomal_protein_S8 | Translation, ribosomal structure and biogenesis | 4486 |
| GBS_RS04880 | gbs0896 | alpha-ketoacid_dehydrogenase_subunit_beta | Energy production and conversion | 4394 |

<sup>a</sup> Open Reading Frames (ORFs) designations according to NEM316 genome (GenBank accession number NC004368)
<sup>b</sup> Predicted function according to new genome annotation for NEM316, updated at 07-JUN-2020 (GenBank version NC004368.1)
<sup>c</sup> Functional categories classified by RPS-BLAST against the COG database
<sup>d</sup> tRNA and rRNA were excluded as their expression levels are biased due to mRNA enrichment procedure.

The functional category “Signal transduction mechanisms” was represented among the top-100 most expressed genes by only one gene, GBS_RS08755, a response regulator transcription factor, CovR, known to play a role on virulence gene expression [29, 30].

Finally, we were interested in assessing the expression levels of genes of interest (n=47) (Figure 2). The set includes the seven genes that putatively encode for secreted DNases (GBS_RS03720, GBS_RS04825, GBS_RS01045, GBS_RS03490, GBS_RS02295, GBS_RS03960, GBS_RS05380) and the genetic pilus islands (PI), that consists of five genes which encode for pilus assembly [31], PI-1 (that plays an important role in evasion from host innate immunity) (GBS_RS03580, GBS_RS03575, GBS_RS03570, GBS_RS03585 and GBS_RS03565) and PI-2a (that is specifically involved in adhesion and biofilm formation) (GBS_RS07760, GBS_RS07745, GBS_RS07765, GBS_RS07755 and GBS_RS07750) [32, 35]. The major nuclease, nuclease A (GBS_RS03720; old locus tag: gbs0661), identified in *S. agalactiae* NEM316 by Derré-Bobillot and co-authors [36], presented higher mean expression values (ranked at position 520° out of a total of 2169 genes evaluated) than the other DNase genes (Figure 2). Among the PI genes, PI-2a presented higher mean expression values (ranked between positions 84° and 718° out of a total of 2169 genes evaluated) than PI-1 coding genes (ranked between positions 1058° and 1973° out of a total of 2169 genes evaluated) (Figure 2).

**Figure 2.**
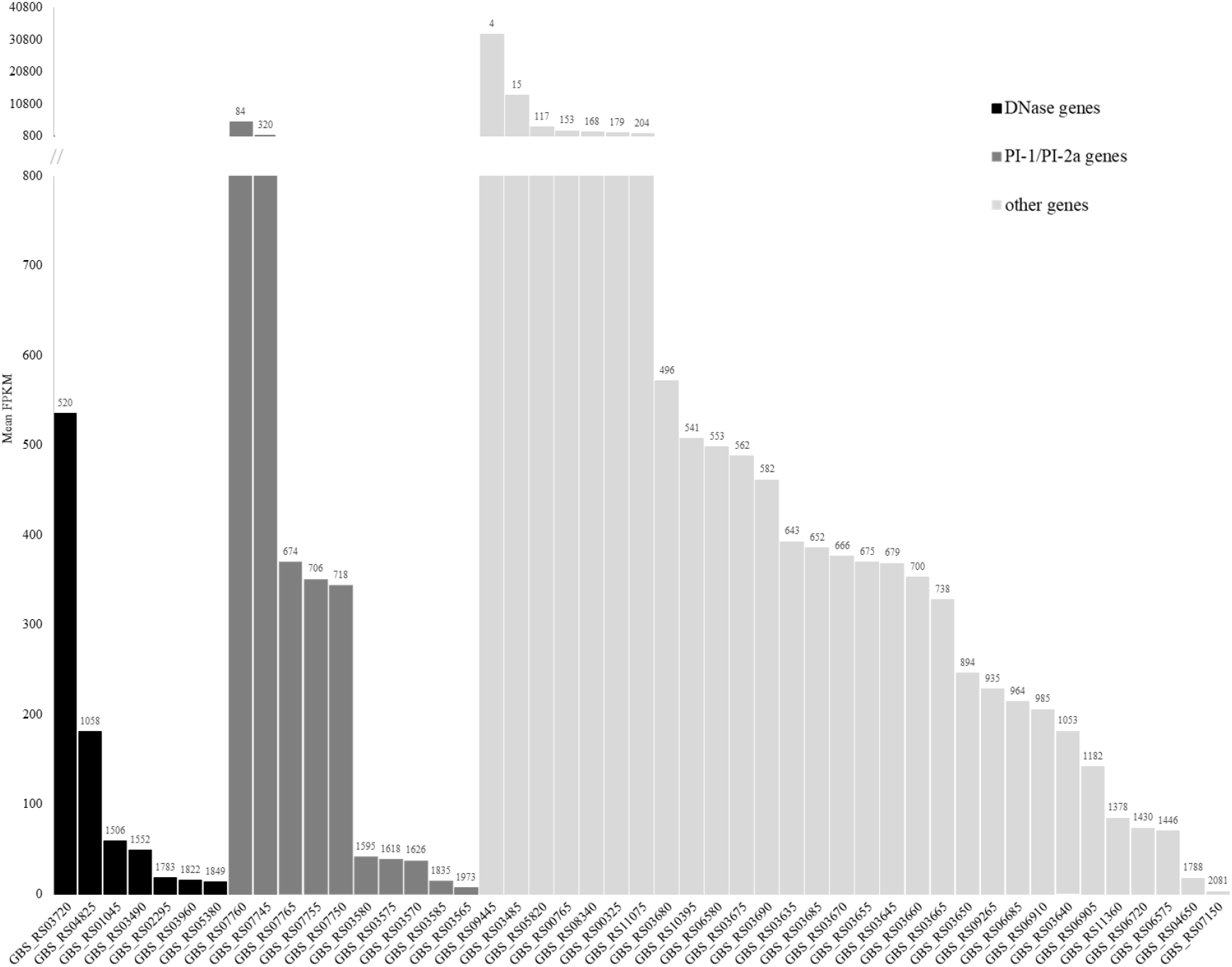
Expression level (FPKM units) of genes potentially involved in virulence in reference strain NEM316 at exponential growth phase. Genes are grouped according to their predictive function in three groups, DNase, PI-1/PI-2a and other coding genes, and displayed by increasing level of expression. The values above the bars correspond to the rank position of each gene in the total of 2169 genes evaluated.

Interestingly, three virulence associated genes were ranked within the top-100 most expressed genes, including the PI-2a pilus major subunit PilB (GBS_RS07760) (Figure 2). The other two genes code for adhesins associated to the glycolytic pathway for energy metabolism; they were GBS_RS09445 (Glyceraldehyde-3-Phosphate Dehydrogenase, GAPDH) and GBS_RS03485 (enolase), belonging to “Carbohydrate transport and metabolism” functional category [37, 38]. GAPDH is also thought to be involved in macromolecular interactions and bacterial pathogenesis, enhancing *S. agalactiae* virulence.

### 3.3. NEM316 comparative transcriptomic analysis through exposure to DNA

The scientific interest in *S. agalactiae* extracellular nucleases increased with the discovery, by Brinkmann and colleagues in 2004 [39], that they can disrupt the DNA matrix, which constitutes the nuclear backbone of neutrophil extracellular traps (NETs). Indeed, as NETs (also composed by granule proteins and histones) [36, 40-41] are released by neutrophils to degrade virulence factors and to kill bacteria, this nuclease-mediated mechanism could play an important role in *S. agalactiae* virulence. Since then, although important knowledge has been acquired about *S. agalactiae* nucleases [4, 36, 42-43] very little is known about the complex molecular pathways and stimuli triggering the release of DNase’s *in vitro* and *in vivo.* Here, we conducted preliminary RNA-seq assays to understand whether direct DNA exposure could affect gene expression during the exponential growth phase of the DNase producer reference strain NEM316. Although a preliminary assay at the transcriptomic level targeting *gbs0661* revealed no significant expression differences among reference strains NEM316 (DNase producer) and 2603V/R (DNase non-producer) with and without DNA stimuli (data not shown), we hypothesized that exposure to human DNA could trigger differential gene expression in other bacterial genes. These genes could potentially be involved in molecular cascades mediating DNase release and virulence. However, this hypothesis was not verified, as no differentially expressed genes were detected for NEM316 during 20 minutes of the exponential phase (at 37°C in THB), either with or without the presence of human DNA (interactive online data navigation is available here: http://degust.erc.monash.edu/degust/compare.html?code=b11b5fff2bf525ea465dc450989f351e#) (Figure 3). While this data could suggest that human DNA, as a stimulus, has no impact on *S. agalactiae* transcription, there is a need to test other stimuli as well as other strains. In fact, DNases have been shown to be under the control of the extensive regulatory systems in streptococci [43] and further work is required to fully understand the complex regulation of the expression of DNases. Thus, it would be interesting to perform comparative RNA-seq assays involving both high and low DNase producing strains in comparison with DNase non-producers. Also, human neutrophils and NETs could possibly be used as stimuli that better mimic the *in vivo* infection environment. On the other hand, testing *S. agalactiae* strains of human clinical origin, from carriage and invasion, might also be of interest. However, we cannot exclude the putative role of DNases in enzymatic kinetics, i.e., in the catalytic mechanism, and in their activity.

**Figure 3.**
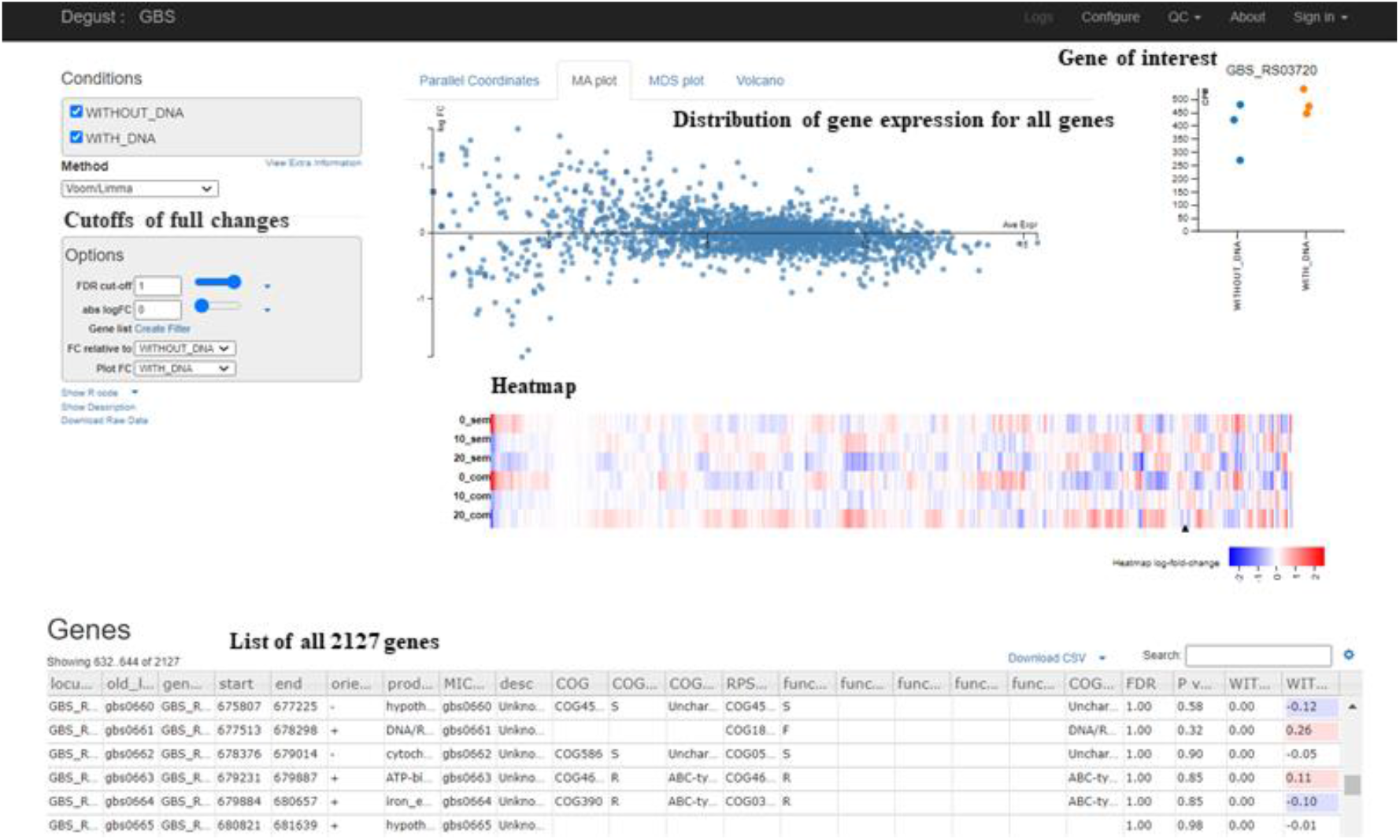
DEGUST RNA-seq database of NEM316 growing at exponential phase with and without DNA exposure. Interactive web tool showing comparative global gene expression analyses between normal growth condition and the “direct DNA exposure” of NEM316 exponentially growing during 20 minutes.

This preliminary assay consolidated our expectation that follow-up studies are needed to disclose the complex molecular pathways (and putative stimuli) triggering the release of DNases. The public release of our data (counts per million, logFC and differential expression statistics, etc) to the scientific community through an interactive and user-friendly web tool (http://degust.erc.monash.edu/degust/compare.html?code=b11b5fff2bf525ea465dc450989f351e#) might further help future analysis and interpretation of other RNA-seq studies in *S. agalactiae*.

## 4. Conclusion

This study constitutes the first attempt to systematize the expression levels of *S. agalactiae* reference strain NEM316 during their normal growth in the laboratory, namely during the exponential growth phase. By providing a comprehensive comparison of gene expression by gene functional category and exploring the levels of expression of genes likely associated with virulence, our data may constitute a database for future functional studies. Preliminary data generated for differential gene expression through DNA exposure suggests the need of further studies regarding the impact of NETs in bacteria.

