## Supplementary Figure S1 for "Global gene expression analysis of *Streptococcus agalactiae* at exponential growth phase"

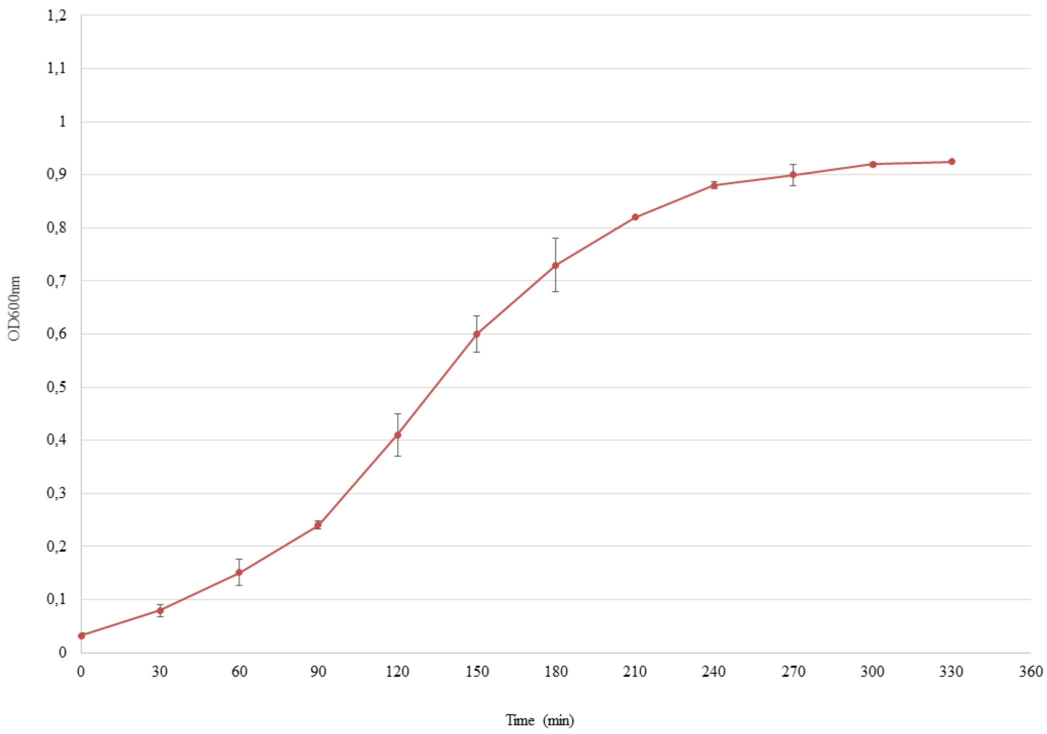


**Supplementary Figure S1 Growth curve of *S. agalactiae* strain NEM316 at 37ºC.** Bacteria were inoculated into THB from an overnight culture and growth was monitored by measuring the $OD_{600}$. Three independent experiments were performed.
