## Supplementary Table S1 for "Global gene expression analysis of *Streptococcus agalactiae* at exponential growth phase"

**Supplementary Table S1** List of genes for which expression values were obtained through RNA-seq for NEM316 at exponential growth phase. Due to the massive extent of this table, it was impossible to present it in a printable format. Please access the following link to review the table: http://doi.org/10.5281/zenodo.4129766
